# Genomic sequences and RNA binding proteins predict RNA splicing kinetics in various single-cell contexts

**DOI:** 10.1101/2021.05.02.442314

**Authors:** Ruiyan Hou, Yuanhua Huang

**Affiliations:** School of Biomedical Sciences, University of Hong Kong, Hong Kong SAR; Department of Statistics and Actuarial Science, University of Hong Kong, Hong Kong SAR

## Abstract

RNA splicing is a key step of gene expression in higher organisms. Accurate quantification of the two-step splicing kinetics is of high interests not only for understanding the regulatory machinery, but also for estimating the RNA velocity in single cells. However, the kinetic rates remain poorly understood due to the intrinsic low content of unspliced RNAs and its stochasticity across contexts. Here, we estimated the relative splicing efficiency across a variety of single-cell RNA-Seq data with scVelo. We further extracted three large feature sets including 92 basic genomic sequence features, 65,536 octamers and 120 RNA binding proteins features and found they are highly predictive to RNA splicing efficiency across multiple tissues on human and mouse. A set of important features have been identified with strong regulatory potentials on splicing efficiency. This predictive power brings promise to reveal the complexity of RNA processing and to enhance the estimation of single-cell RNA velocity.

## Introduction

RNA metabolism is a fundamental process of gene expression which includes RNA synthesis, splicing and degradation. The rates of both RNA splicing and degradation determine the amount of mature RNAs in steady state and the speeds of responding extra- and intra- cellular signals [1]. Therefore, it remains of high interests to accurately measure the kinetic rates in various tissues and conditions. One of the most appealing strategies is to metabolically label and enrich nascent transcripts, such as 4sU-seq [2] and TT-seq [3]. Similar approaches on nascent RNAs labelling have also been extended to single-cell resolution, e.g. scSLAM, scNT-seq and RNA timestamps [4–6], for fully capturing the stochasticity of transcription processes in a heterogeneous cell population.

Additionally, the commonly used RNA-Seq of total or Poly-A RNAs (without metabolic labelling) has also been reported to contain a non-negligible proportion of intronic reads in both bulk [7, 8] and single-cell samples [9]. Increasing evidence suggests that such intronic reads represent the level of pre-mRNAs hence are informative for revealing transcriptional regulation [8, 10] and RNA kinetic parameters [9, 11]. One prominent use of these intronic reads is to estimate the kinetic rates in single cells, followed by quantification of RNA velocity - the time derivative of the mature RNAs, which greatly aids to infer future state of each cell on a time scale of hours [9]. A likelihood-based dynamical model, scVelo, has further enabled the estimation of transcription, splicing and degradation rates by relaxing the observation of full stationary states, hence increasing the applicability [12]. Therefore, RNA velocity methods bring new opportunities to examine splicing kinetics in various fresh tissues and understand their variability that was largely understudied, partly due to the laborious metabolic labelling techniques [13, 14].

Previous observations suggest that the kinetics of RNA metabolism may be determined by genomic sequence elements. The link between intron length and splicing rate has been particularly intensively studied. In *Drosophila melanogaster*, the relationship between intron lengths and the estimated splicing half-lives was overall weak, but the half-life decreases notably when the intron length is shorter [15]. Spliced introns tend to be shorter than unspliced introns in human K562 dataset, which illustrates the importance of intron lengths on splicing from other aspects [16]. Intron secondary structure also contributes to the kinetic of pre-mRNA splicing in yeast and RNAs tends to be spliced faster when their introns have a less favorable predicted secondary structure [17]. Additionally, the increased motif strengths of the two splice sites (5’ss and 3’ss) were proved to be related with shorter intron half-life [15]. A single-nucleotide deviation from the consensus motif on splice site and branchpoint can change synthesis time or donor bond half-life [14]. Furthermore, the octamers features also play an important role in predicting donor-bond and acceptor-bond half-life [14], presumably in cooperation with RNA binding proteins (RBP) [18]. However, little is known about the relationship between these features and RNA splicing kinetics at single cell resolution, and how predictive of these integrative features across different tissues.

In this work, we asked whether these sequence features can predict splicing efficiency measured from single-cell RNA-seq (scRNA-seq) data. We first estimated the splicing rate and degradation rate by using the stochastic model from the scVelo package in K562 cell and defined the relative splicing efficiency as the ratio of splicing rate and degradation rate. Then we built three feature sets including the 92 basic features, the RNA binding protein (RBP) features and the octamer features (Fig. 1; details below). Next the seven combinations of these three feature sets were used to predict the relative splicing efficiency by fitting random forest regression models. In addition, we also evaluated the importance of different features and identified a set of determinant factors for the relative splicing efficiency. To investigate the diversity of the splicing efficiency and the wide applicability of our model, we collected datasets from five tissues in human or mouse and performed both within and cross tissue predictions. Finally, to prove our model’s reliability in different technology, we leveraged our curated features to predict splicing efficiency quantified from other assays.

**Figure 1.**
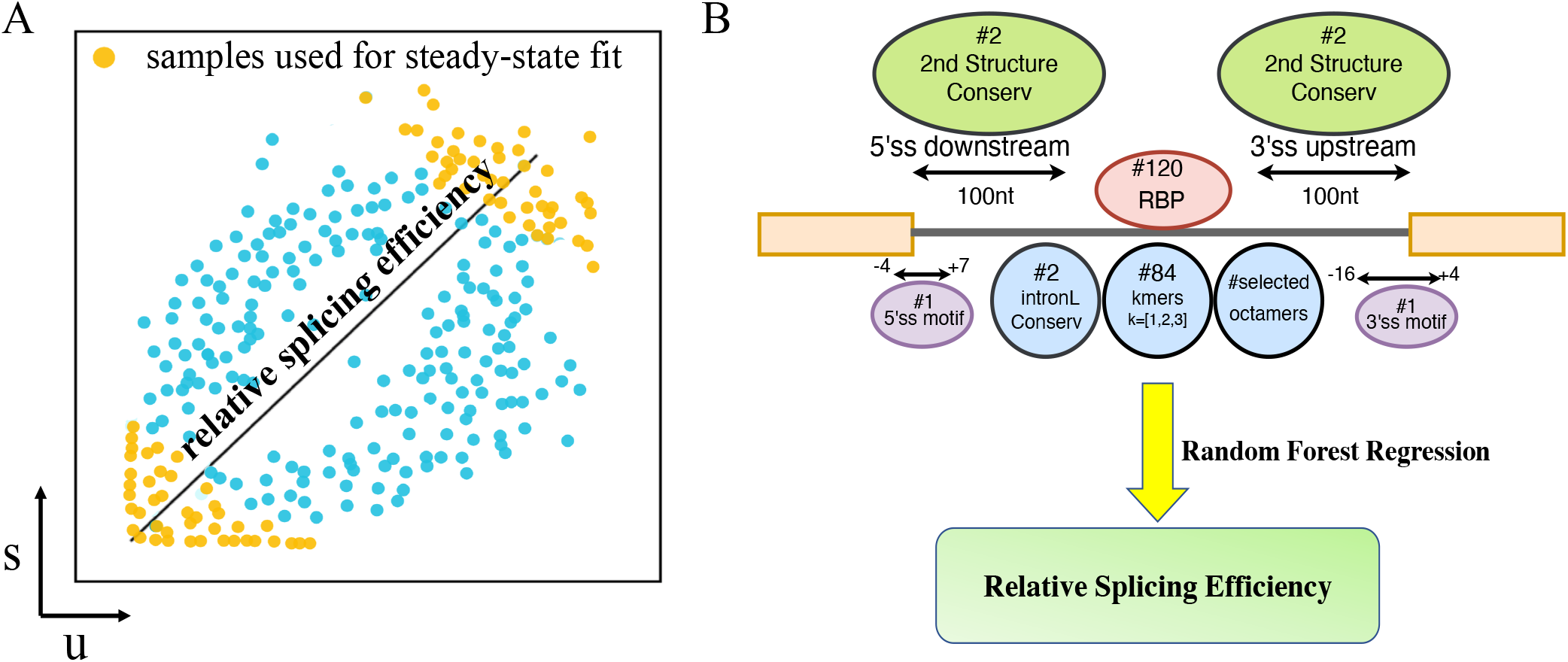
Quantification of relative splicing efficiency from scRNA-seq data and its prediction with genomic related features. **(A)** The relative splicing efficiency is defined as the ratio of the splicing rate and degradation rate, which can be quantified through the spliced and unspliced read counts of cells in steady states. X-axis and y-axis are the normalised read counts for unspliced (U) and spliced (S) RNAs, respectively. Each dot is a cell, with gold color denoting cells in steady states. **(B)** A schematic of predicting relative splicing efficiency with three genomic feature sets: 92 basic sequence features, 120 RBPs features and variable number of pre-selected octamers. Random forest regression model is used for the prediction by default.

## Results

### Splicing efficiency quantified with RNA velocity method

In the work, we employed a recently proposed RNA velocity method scVelo to estimate the splicing kinetic rates in single-cell RNA-seq (scRNA-seq) data. Briefly, this method measures the abundance of unspliced (U) and spliced (S) RNAs and it assumes that under a linear kinetic model the derivative of S, d*S*/d*t* = *βU* − *γS*, is 0 for cells in steady state, where *β* and *γ* denoting for splicing and degradation rates, respectively. Therefore it quantifies the ratio between U and S across a group of cells in steady state as an estimate of the relative degradation rate. Conversely, its reciprocal is the relative splicing rate (or splicing efficiency, i.e., the ratio of splicing rate and degradation rate) that can be approximated by the ratio (slop) between S and U (Fig. 1A). Besides the steady-state setting in stochastic mode, scVelo also provides dynamical mode by relaxing the requirement of observing two stationary states.

We first explored the dynamical and stochastic modes in scVelo and found that the steady state-ratio (degradation versus splicing rate) in both modes are highly correlated (Pearson’s R=0.69 to 0.98 in six data sets; Supp. Fig. S1). Interestingly, the decoupling of the splicing and degradation rates in the dynamical mode retains the major variability in splicing rate, which explains 48.8-96.9% of the variance on relative splicing efficiency measured in the stochastic mode (Supp. Fig. S2b; with variancePartition package [19]) and consequently their high correlations (Supp. Fig. S2a). Therefore, we quantify the (relative) splicing efficiency via the stochastic mode in scVelo for its simpler model assumptions. Unsurprisingly, we found that this quantification is highly consistent with two other tools: velocyto [9], another RNA velocity method with similar settings, and INSPEcT- [11], an RNA kinetics estimator with treating scRNA-seq data in a bulk manner (Pearson’s R=0.97 for velocyto and Pearson’s R=0.667 for INSPEcT-; Supp. Fig. S3).

With the above quantification, we first examined six scRNA-seq data sets on mouse and human, and found that relative splicing efficiency generally displays a unimodal Gaussian-like pattern at logarithm scale, while the mean varies across tissues, e.g., mouse lung has lower splicing efficiency compared to mouse pancreas (Supp. Fig. S4).

### Genomic features accurately predict splicing kinetics

Next, we introduced three sets of genomic features extracted from the protein coding genes and their introns in human and mouse to predict the relative splicing efficiency (Fig. 1B and Supp. Table S1). Specifically, in our first feature set, we have defined 92 sequence related features, including intron length, two RNA secondary structures (Delta G) for the region 100nt downstream of 5’SS and the region 100nt upstream of 3’SS, three conservation scores for the above regions and intron, two motif scores for the 5’SS and 3’SS and the frequency of 84 kmers (K=1, 2, 3). Secondly, we calculated the frequency of 65,536 octamers in the whole intron, which are further selected by a Lasso regression model, resulting in 36-121 features left in different data sets (Supp. Fig. S5; Methods). Last, we also included 120 RNA binding proteins, and used their density on the whole gene body as our third feature set, though it is only available for K562 cell line [20]. Note, we took each gene as a sample by either measuring features at gene level (RBPs), or aggregating intron-level features per gene with median value for 92 basic features and mean for octamers.

We then leveraged the above defined the genomic features to predict the splicing efficiency in K562 cell line and found it can be accurately predicted by a random forest model (Pearson’s R=0.754, Fig. 2A). We noticed linear models, including Lasso with sparse weights, do not work well (Supp. Fig. S6), suggesting non-linear relationship between features and splicing efficiency, partly due to the wide existence of zero-count features.

**Figure 2.**
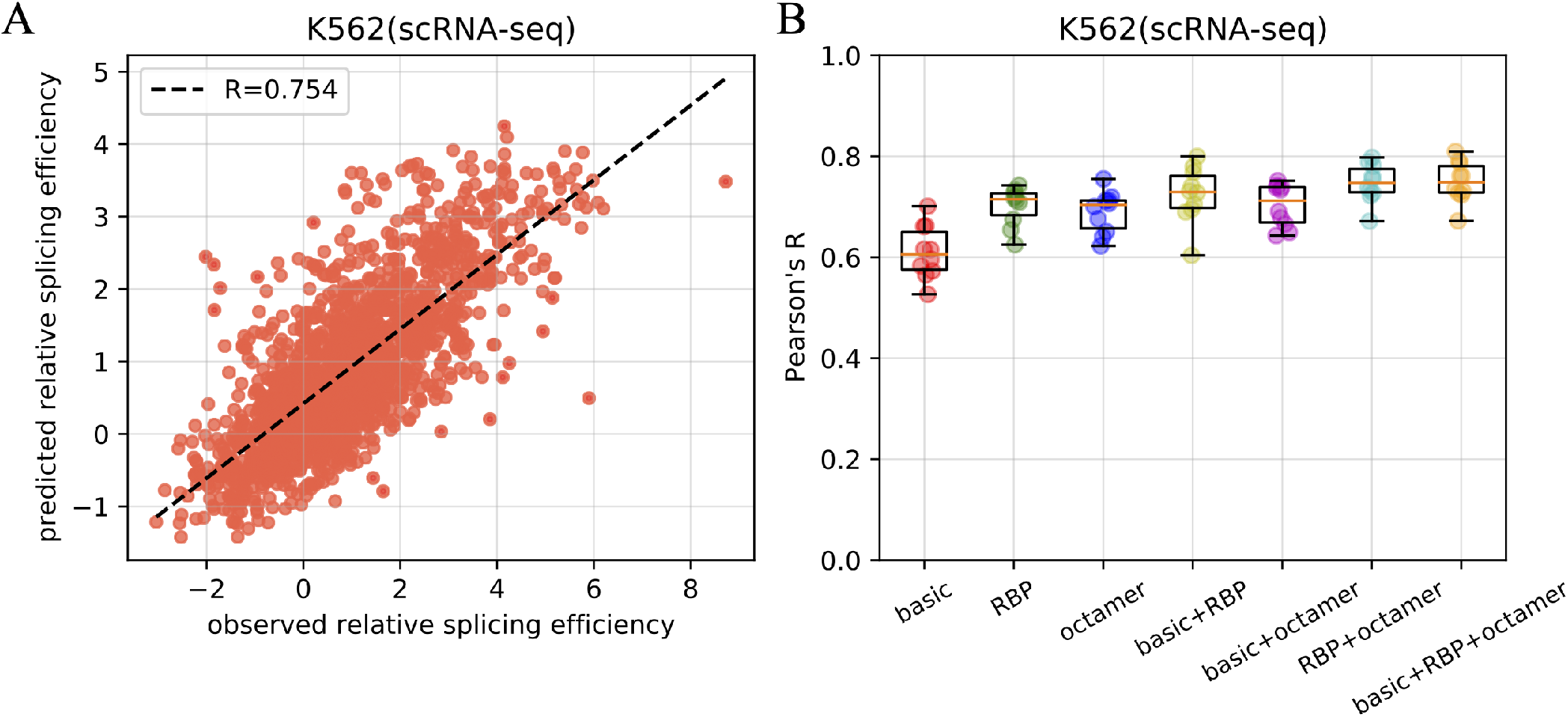
The splicing efficiency can be accurately predicted by genomic features. **(A)** A scatter plot between the predicted and measured relative splicing efficiency from K562 scRNA-seq data with full genomic features. **(B)** Three genomic feature sets and their combinations are used to predict the splicing efficiency in K562 cells as (A). Random forest regression models are applied here, and predicted values are obtained via 10-fold cross validations.

To further confirm the reliability of our model, we examined a null setting by permuting genes and found the regression model cannot predict non-correspondent splicing efficiency at all (Pearson’s R=0.058; Fig. S7). Within these three features sets, Fig. 2B shows the RBP features are the most predictive one (Pearson’s R=0.702) on predicting splicing efficiency, followed by octamer features set (Pearson’s R=0.690) and 92 basic features set (Pearson’s R=0.610). Additionally, the combinations between two feature sets perform better than any set alone, and the integration of full features achieves the best performance, which suggests each feature set contains independent prediction power.

### Feature importance and cooperative structure

Next, we further studied how each specific feature contributes to the prediction of splicing efficiency. In the full feature set model, octamers and RBPs dominate the top features, as expected from the most predictive performance. Specifically, PABPC4 is the most important features (Fig. 3A). It has been reported that PABPC4 prefers to bind polyadenylate [21] and was associated with an increase splicing efficiency (Supp. Fig. S8 and Supp. Table S2). AAAGAAAA is one of the most important octamer features (Fig. 3A and Supp. Fig. S9). We speculate the reason is that AAAGAAAA motif can be bound by SRSF4 [22], which is associated with the splicing of detained introns (DI) and this step may slow down the splicing process [23]. The serine/arginine-rich proteins were reported to regulate the donor and acceptor bond half-life in different directions depending on their binding sites [14].

**Figure 3.**
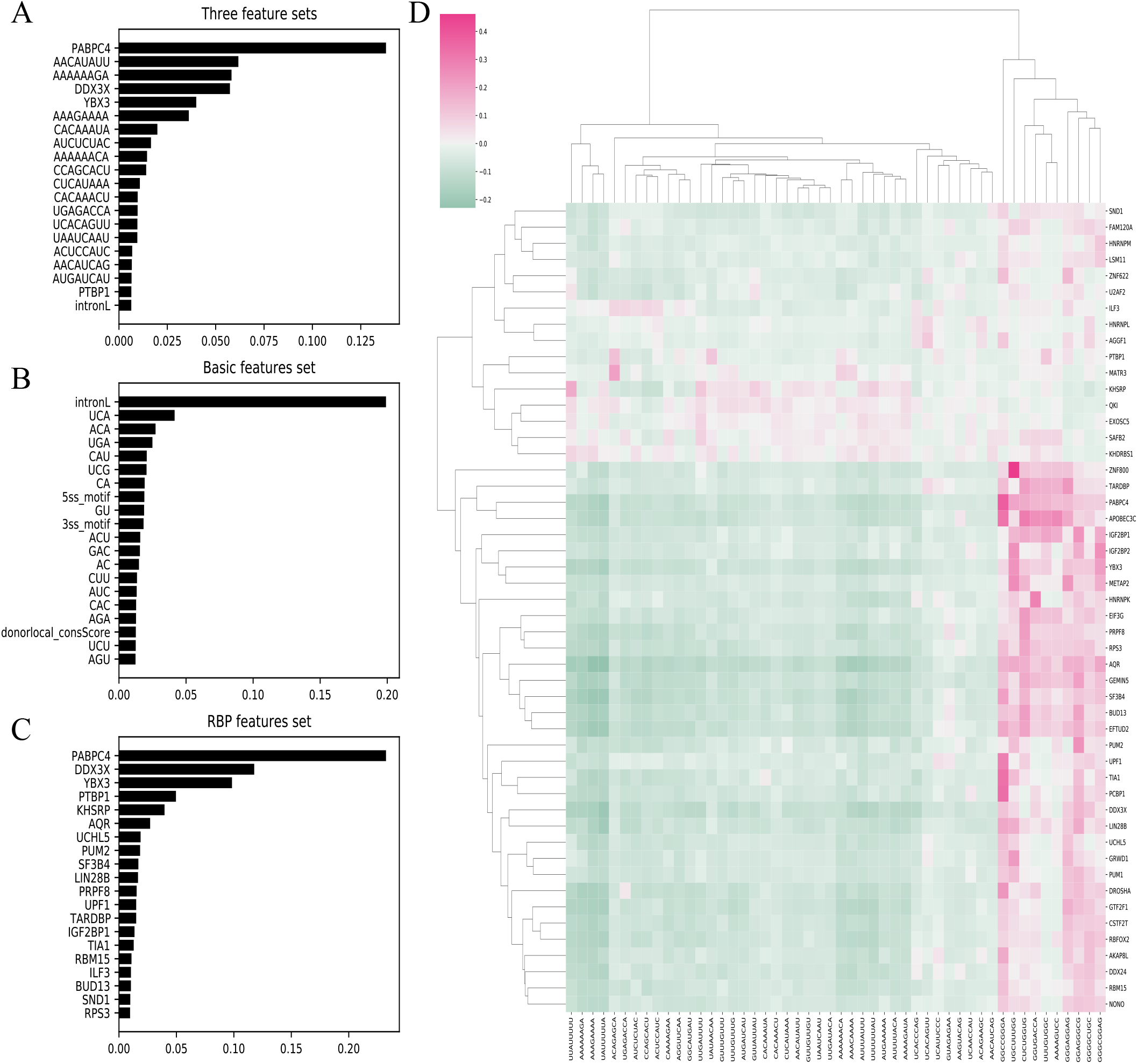
The top important features in different feature sets and the relationship between the octamers and the RNA binding proteins. The top 20 important features when using full features to predict: the 92 basic features, octamer features and 120 RBP features **(A)**, or only the 92 basic features **(B)**, or only the 120 RBP features **(C)**. **(D)** The correlation of top 50 octamers’ counts and top 50 RNA binding proteins’ densities.

Furthermore, we zoomed into the prediction model with only one individual feature set and evaluated the feature importance in the prediction. Among the 92 sequence related features, the importance of intron length is much higher than any other feature (Fig. 3B). Remarkably, there was a negative correlation between the intron length and splicing efficiency (Fig. S8). This result showed that short introns have faster splicing rate as in agreement with the observation in mouse [1]. However, this is in contrast to the observation in the *Schizosaccharomyces*, where the intron length did not correlate with the splicing time [24]. This could be due to the closer evolutionary relationship between human and mouse. The frequency of UCA also plays a vital role in splicing efficiency (Fig. 3B). This result is in agreement with previous publication that UCA shows significantly high correlation with the splicing speed in non-RP (ribosomal protein) genes in yeast [17].

In RNA binding proteins, high importance indexes were observed at PABPC4, DDX3X, YBX3 and PTBP1 (Fig. 3C). DDX3X participates in all the stage of RNA metabolism, and functions as an essential component of messenger ribonucleoproteins, human spliceosomes and spliceosomal B complexes [25–27]. DDX3X positively correlates with the relative splicing efficiency, likely because DDX3X affects the rate of spliceosome assembly. Several publications have already shown that YBX3 plays an important role in translation and mRNA stability [28]. PTBP1 also plays a vital role in splicing efficiency, in agreement with a previous report showing that PTBP1 regulates donor bond half-life in positive or negative fashions [14]. Interestingly, A-rich octamers in intron region significantly influence the splicing efficiency (Supp. Fig. S9) and the reason need to be further elucidated.

We further assessed the correlation structure of a subset of predictive octamer and RBPs, and noticed that ZNF800 has the strongest positive correlation with the UGCUUUGG. Additionally, there was a negative correlation between the AAAAAAGA and most of RBPs, which may explain how it contributes to the regulation of splicing efficiency (Fig. 3D). Also, as the most predictive feature, PABPC4 positively correlates with a range of octamers, specially GGCCGGGA.

### Consistent kinetic regulations across tissue and species

Next, we ask whether the genomic regulatory pattern can also be observed in other tissues and species. We further applied scVelo to five more data sets, including PBMC and lung in human, and dentate gyrus, pancreas and lung in mouse. As shown in Supp. Fig. S10, the relative splicing efficiency is only moderately correlated between tissues and species (Pearson’s R from 0.306 between PBMC and DG to 0.706 between mouse lung and pancreas). The intra-species pairs shows are generally higher correlation between the inter-species pairs, probably due to evolutionary divergence.

Then we used our model to predict the splicing efficiency on each of the five tissues. Because the number of octamer features selected by Lasso are different across tissues, we took the octamer features selected on K562 as the general set for all other tissues with the premise that the prediction performance largely remains with octamers selected from its own data set or K562 (see human lung dataset as example in Supp. Fig. S11). The full feature set, including the selected octamer features (d=96), the 92 related to sequence features and the 120 RBP features, were applied to predict the splicing efficiency in these tissues mentioned above. The strong predictive power of our model was confirmed by high Pearson’s R, ranging from 0.500 to 0.771, between the predicted and measured relative splicing efficiency (Fig. 4A).

**Figure 4.**
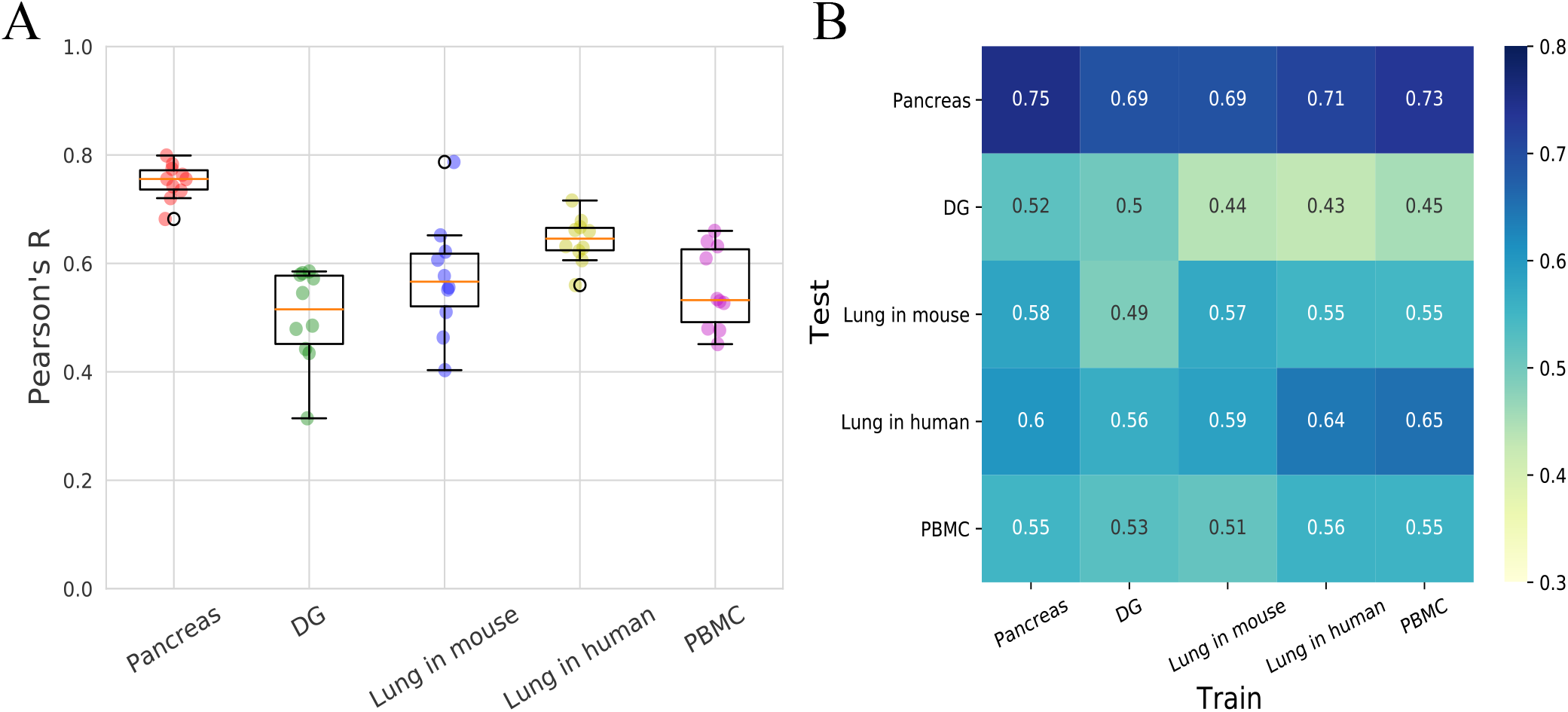
The relative splicing efficiency can be predicted across different tissues. **(A)** Prediction performance in Pearson’s R between predicted and measured relative splicing efficiency on five tissues. Each dot denotes a fold in ten-fold cross-validation. Random forest model is used here with the integrative features: basic features set, RBP features set and octamer features set. **(B)** Prediction performance for within tissue (diagonal) and cross-tissue (off diagonal) prediction. Shown is Pearson’s R as in (A).

Furthermore, we evaluate the extent to which the regulatory of kinetic rates are shared across tissues and species. To address this question, we fit the regression model from one tissue and predict the splicing efficiency on another tissue. We found that the cross-tissue prediction is highly consistent with the within-tissue prediction via cross-validation (difference within 0.02 between cross-validation and the best cross-tissue setting; Fig. 4B). On the other hand, the variability across test tissues is dramatically higher (column-wise std: 0.072-0.096) than across training tissues (row-wise std: 0.018-0.035). In other words, if a tissue is less predictable by itself e.g., DG or lung in mouse, it cannot be significantly improved by models fit from other tissues (Fig. 4B).

Taken together, the splicing efficiency is generally predictable across multiple tissues in mouse and human and their regulatory patterns are largely shared. Nonetheless, we noticed some tissues may have lower prediction performance, presumably due to higher noise in the estimation of splicing efficiency or increased complexity of regulation beyond genomic features.

## Discussion

RNA splicing is fundamental to the adjustment of gene expression, and deciphering its precise regulation has been a central focus in molecular biology for the past few decades. Very recently, the RNA splicing kinetics have drawn great attention again, thanks to the emergency of RNA velocity in single cells that leverages the intrinsic RNA processing dynamics as footprints for inferring the transient cellular differentiation [9]. Both computational tools [9, 12, 29, 30] and experimental technologies [4–6] have been rapidly developed to more accurately estimate the splicing kinetic rates and / or the cell transitions on top of it. On the other hand, due to minutes of the initial RNA contents, a subset of genes may suffer from higher technical variability in measuring splicing efficiency and consequently affects the quantification of RNA velocity, as the (relative) splicing rate functions as a key parameter there. Therefore, the accurate predictions that we achieved here with genomic features could largely relieve this bottleneck challenge by adaptively imputing the splicing efficiency in a specific tissue or condition.

Though scRNA-seq and the RNA velocity framework opens a new venue to estimate splicing kinetics, it remains challenging to obtain the absolute values of synthesis, splicing and degradation rates. This is because the observations of spliced and unspliced reads are cooperative results from kinetic rates and its latent induction or suppression time [12]. On the other hand, the relative splicing efficiency, i.e., the ratio between splicing and degradation rates, is more straightforward to obtain, with proportional to the spliced RNAs [11]. This relative value is also an important indicator of the RNA processes and a key element of the RNA velocity analysis. Additionally, the quantification of relative splicing efficiency from scRNA-seq has a good match between that from bulk RNA-Seq by INSPEcT- (Pearson’s R=0.411, Supp. Fig. S12), considering both quantifications have a certain level of uncertainty. Though different assays may have their own technical effects and noises on (relative) splicing rate measurement, we found that our extensive genomic features also achieve high prediction performance in other quantifications, including from bulk sequencing data with or without metabolic labelling (Pearson’s R=0.532 & 0.501, Supp. Fig. S13).

Given the convenient access to splicing efficiency from common scRNA-seq data, there are opportunities to further extensively examine the variability of splicing rates in different tissues, conditions and cellular states. From this point of view, it also demands more sophisticated models to jointly analyse multiple datasets and understand the precise control of splicing processes in a global or gene-specific manner. More broadly, the genomic feature sets we curated here could function as a pivot to study the genetic effects on splicing rate in a population scale and to investigate the impacts of somatic mutations on dysregulation of splicing processes and disease association.

Taken together, the relation revealed in this study between splicing efficiency and genomic features, including intronic sequences and RNA binding proteins, offers a detailed picture of the regulation of splicing kinetics. The accurate prediction of splicing rates will further enhance the quantification of RNA velocity and broaden its applicability.

## Materials and Methods

### Data

The scRNA-seq dataset containing K562 was downloaded from the SRA database (SRR7646179; SRR7646180) and then demultiplexed by k-means from Cellranger v3.1 to isolate K562 cells [31]. The datasets of pancreas and dentate gyrus are the builtin sample datasets in scvelo package and can be used directly via the function scvelo.datasets [12]. The dataset of 1k PBMC was downloaded from 10× Genomics website (https://support.10xgenomics.com/single-cell-gene-expression/datasets/3.0.0/pbmc_1k_v3?) and obtained by 10× Genomics v3 chemistry platform. The two datasets about lung tissue in mouse and human are from a benchmark study which is based on 10× Genomics platform (v2 kit) [32], with four libraries belonging in to two samples (GEO: GSE133747). The dataset of K562 sample sequenced by TT-seq was downloaded from the GEO data set GSE129635 [14]. Bulk RNA-seq library for K562 cell line comes from the ENCODE project and can be downloaded from the SRA database (SRR4422447; SRR4422446).

### Relative splicing efficiency estimation

The raw fastq files were mapped to human reference genome hg38 and mouse reference genome mm10 by using CellRanger v3.1, respectively. Then we used velocyto [9] to annotate the spliced and unspliced reads. The loom file was obtained and then put into scvelo which is the tool to calculate the splicing rate and degradation rate. The stochastic mode was used to estimate these values. Data was filtered by setting the parameter min shared counts as 20 and n top genes as 2000 in all datasets. The n pcs was set as 30 and n neighbor was set as 30 when computing moments. The relative splicing efficiency was defined as the ratio of splicing rate and degradation rate. For the dataset of K562 (TT-seq), the relative splicing efficiency was calculated through mean of donor bond half-life and acceptor bond half-life divided by junction bond half-life.

### Feature extraction

There were three feature sets generated at intron and gene level in human and mouse. The intron length, splice site motif strength and kmers (k=1,2,3) were extracted by pyseqlib v0.2 [13]. BigWigSummary was used to extract conservation scores for the sequences of 5’ss downstream 100nt, 3’ss upstream 100nt and intron from hg38.phastCons100way.bw or mm10.60way.phastCons.bw [33] for human and mouse, respectively. RNAfold in ViennaRNA v2.4.14 [34] was used to produce secondary structure of 5’ss downstream 100nt and 3’ss upstream 100nt. The frequency of octamer in intron sequence were obtained by the same pyseqlib package. Then the glmnet python was applied for octamer selection with lasso regression and the optimal values of λ for weight penalty was chosen at which the minimal MSE is achieved by using 3-fold cross-validation. Then we filtered those octamers whose coefficients are 0. The largest eCLIP dataset including RNA sequence that combined with 120 RBP in k562 cells was downloaded [20] and BedTools 2.29.2 [35] was executed to intersect the eCLIP peaks and whole gene body, setting that the least overlapping parameter as 50%. The sum of density of all positions in every RBP file were regarded as different RBP features, leading to 120 RBP features in total. The RBP features are directly measured at per gene level. The 92 basic sequence features and octamers are measured at intron level, and then aggregated to a gene level by taking the median value (92 basic features) or mean (octamers).

### Splicing rate and degradation rate prediction

The random forest regression, lasso regression and linear regression models all implemented in scikit-learn were employed to predict the log(1/*γ*) in K562 datasets from scRNA-seq and TT-seq, pancreas, dentate gyrus, PBMC, lung tissue in mouse and human data sets. Seven feature groups were obtained by combining three feature sets mentioned above and then put into the random forest regression models. Three-fold cross-validation was used in the prediction process. The Pearson’s correlation coefficients between these predicted and measured values were calculated and displayed.

### Different tissues cross prediction

In order to explore whether the splicing kinetics vary in different tissues and whether the model fit from one tissue can predict another tissue’s splicing efficiency, we took one of the tissues as training set and the remaining tissues as test set. For the within and cross-tissue predictions, the octamer features are all selected from K562 dataset (scRNA-seq), and the RBP features are also borrowed from K562 cells as an approximate.

### Reproducibility and processed data

In order to ensure the reproducibility, all the scripts used for the prediction and figure generation are publicly available at https://github.com/StatBiomed/scRNA-kinetics-prediction. Additionally, all the processed data sets mentioned above are also compiled and available in this repository, including genomic features, RNA binding proteins, and measured and predicted splicing efficiency.

