## Supplementary Figures for "Genomic sequences and RNA binding proteins predict RNA splicing kinetics in various single-cell contexts"

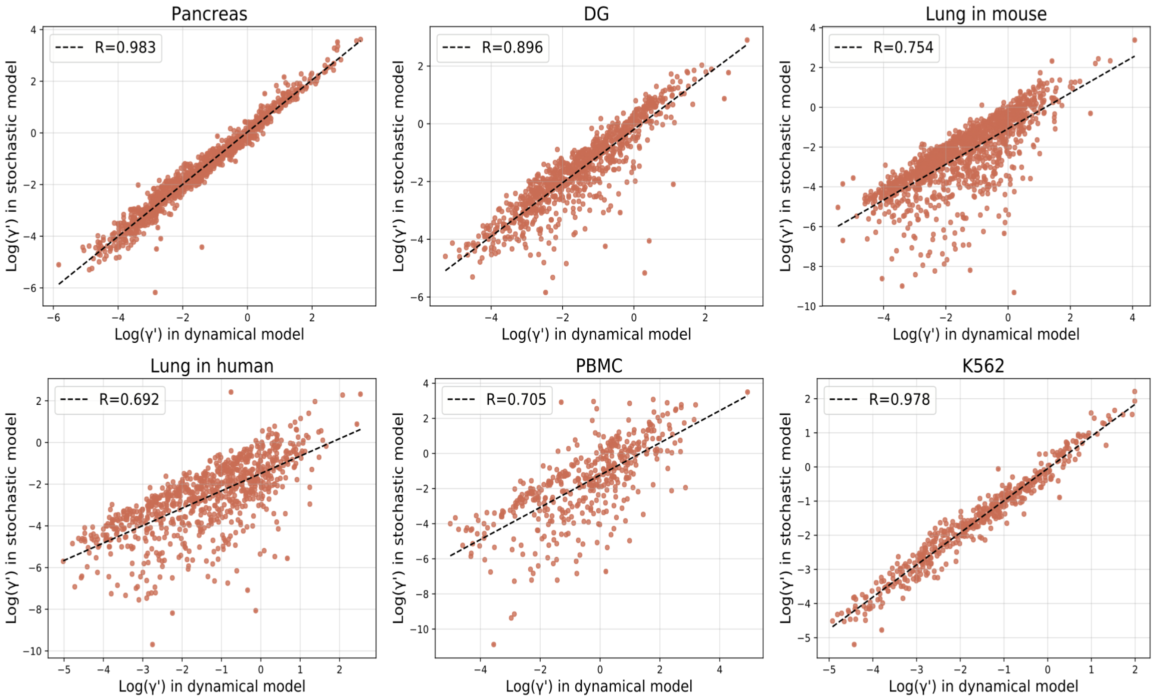


Figure S1: The steady-state ratio correlation between the dynamical model and stochastic model in six scRNA-seq datasets including pancreas, DG, lung in mouse, lung in human, PBMC and K562 datasets. The steady-state ratio ($\gamma'$) is calculated through the degradation rate ($\gamma$) divided by splicing rate ($\beta$). The steady-state ratio between these two models are highly correlated with Pearson's correlation coefficients ranging from 0.705 to 0.983.

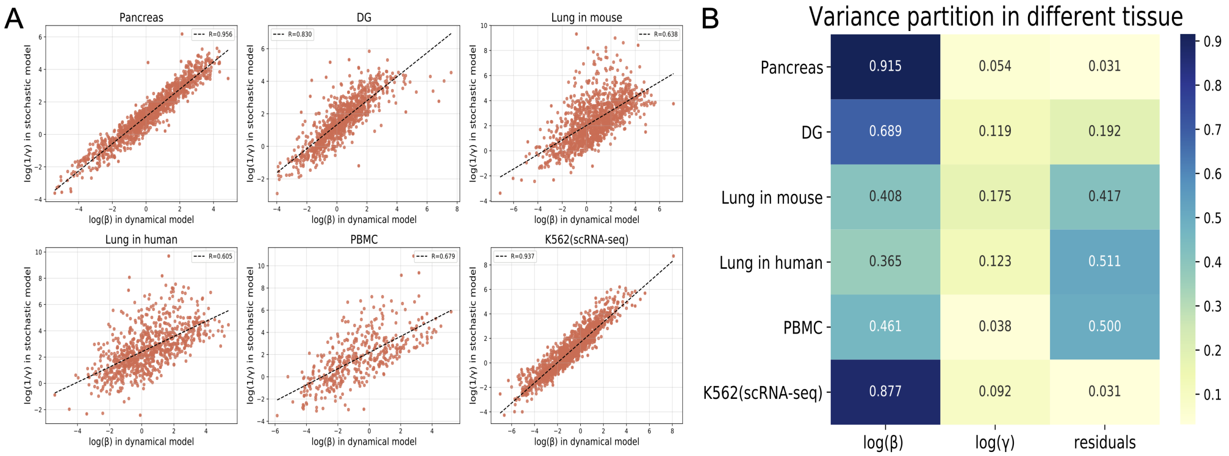


Figure S2: The relationship between the splicing efficiency in stochastic model and the splicing/degradation rate in dynamical model. (A) The correlation between splicing rate($\beta$) estimated by the dynamical model in scVelo and the reciprocal of degradation rate($1/\gamma$) estimated by the stochastic model of scVelo. The Pearson's correlation coefficients are high in different tissues, ranging from 0.605 to 0.956. (B) The contribution of $\log(\beta)$ and $\log(\gamma)$ estimated by dynamical model to variance in the splicing efficiency $\log(1/\gamma_{stoch})$ in stochastic model. The variance composition was calculated through $\log(1/\gamma_{stoch}) \sim \log(\beta) + \log(\gamma) + 1$ by the variancePartition R package v3.2 [1].

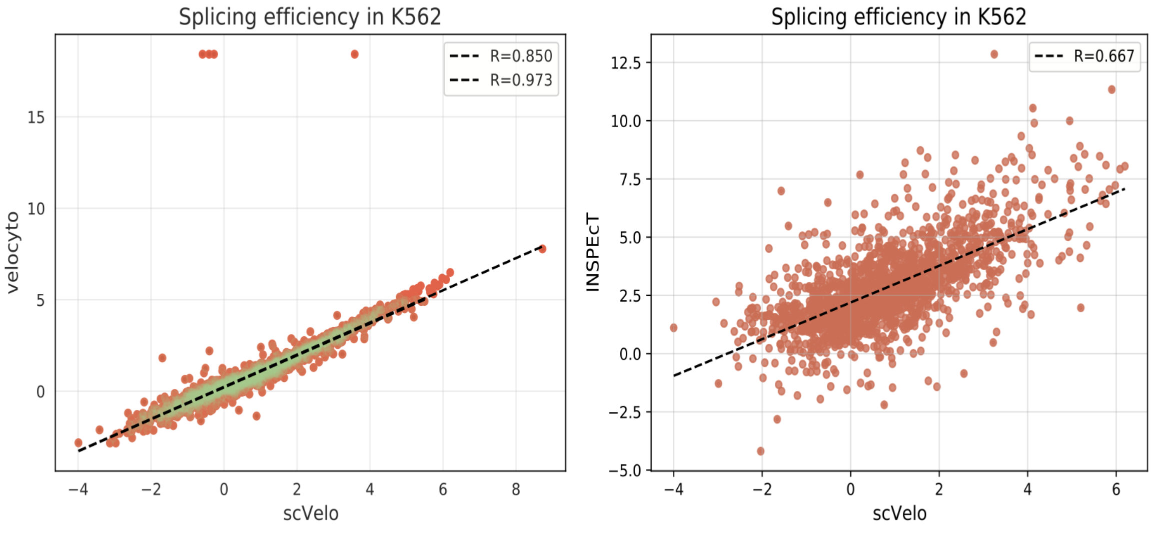


Figure S3: Comparison of splicing efficiency measure between scVelo and other methods. Left panel: scatter plot of log(relative splicing efficiency) in K562 cell line (scRNA-seq) estimated by scVelo and velocyto. The dots in tomato color are all samples (Pearson's R=0.850) and the dots in lime color are samples removing 1% outliers (Pearson's R=0.973). Right panel: scatter plot of log(the relative splicing efficiency) in K562 cell line(scRNA-seq) estimated by scVelo and INSPEcT-.

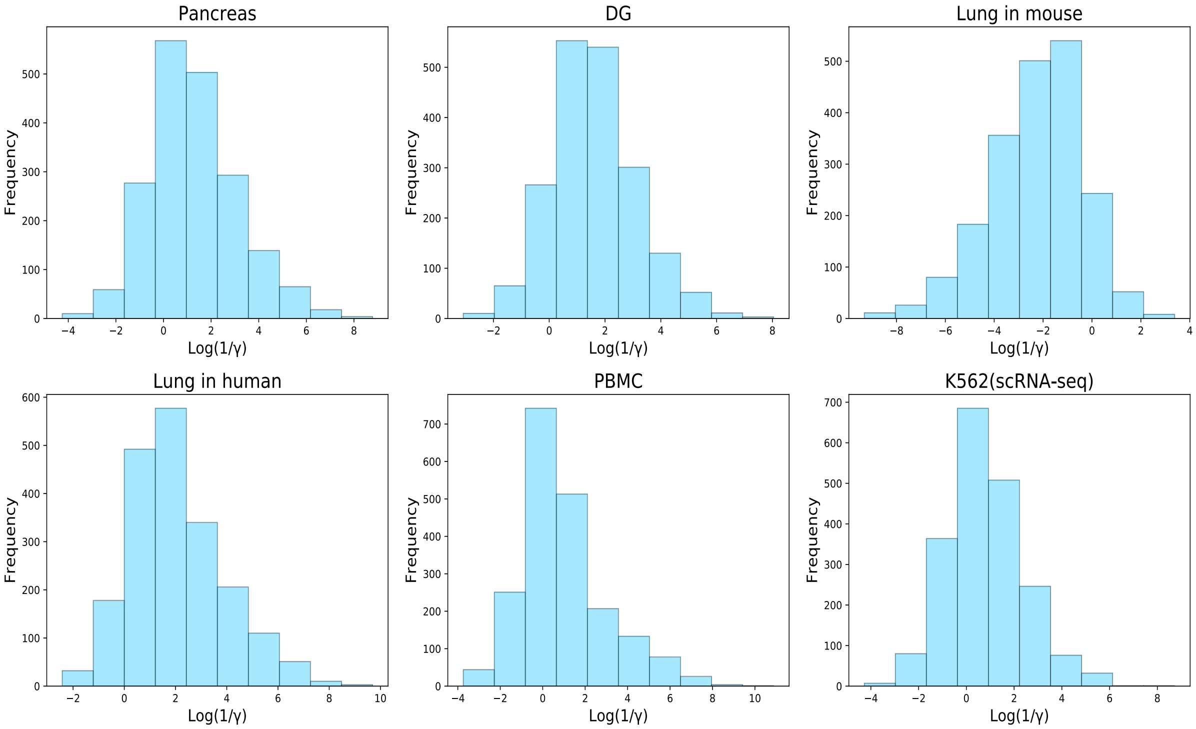


Figure S4: The distribution of the relative splicing efficiency in six scRNA-seq datasets including pancreas, DG, lung in mouse, lung in human, PBMC and K562 datasets. The relative splicing is represented by the ratio of splicing rate and degradation rate. The stochastic model in scVelo is used here.

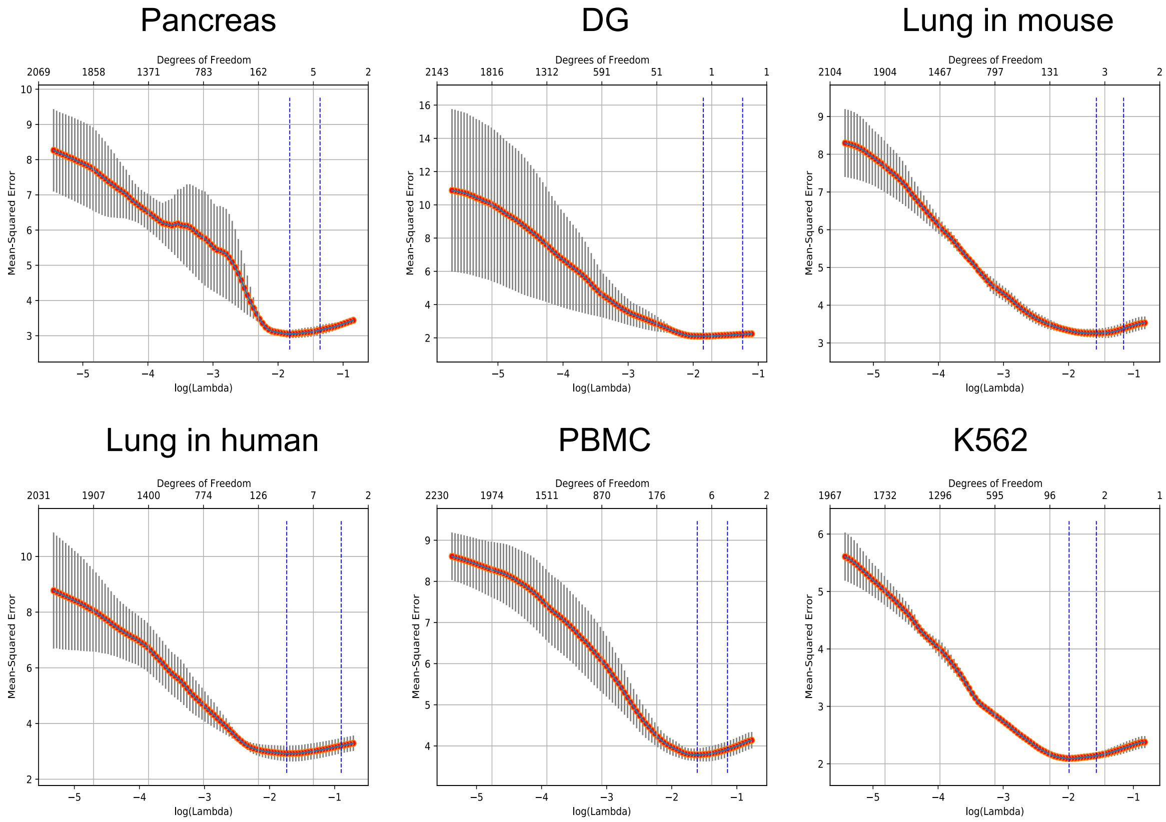


Figure S5: Cross-validation (CV) plot of the Mean-Squared Error (MSE) corrspinding to Lambda by using the python-glmnet (see Methods). 36 to 121 octamers were selected by the lasso regression from 65,536 octamers in different datasets and the number of octamers were determined when the MSE is the smallest.

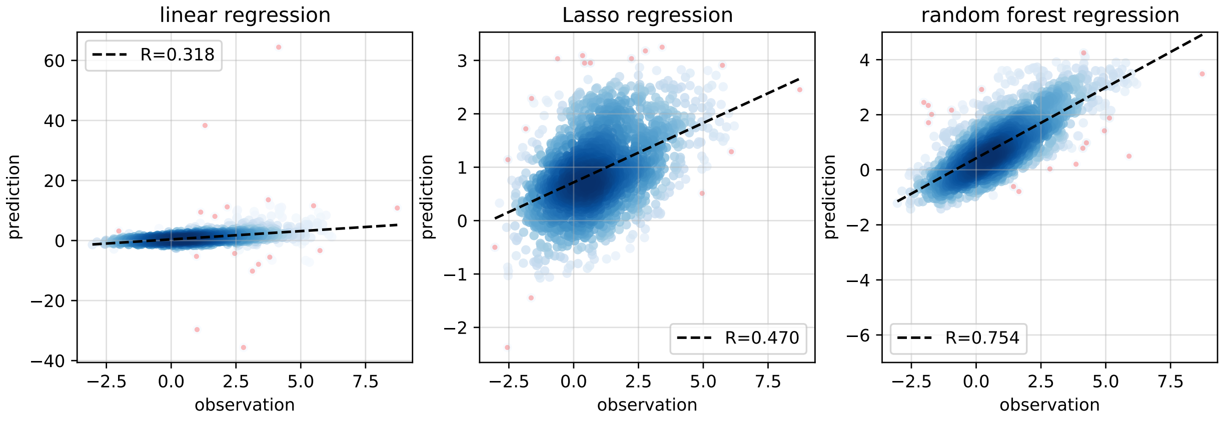


Figure S6: Sequence and RBP features are predictive for the relative splicing efficiency. 92 basic features, 120 RBP features and selected 96 octamers features are applied to predict the relative splicing rate of K562 cell with ordinary linear regression (left panel), Lasso regression (middle panel), and random forest regression (right panel).

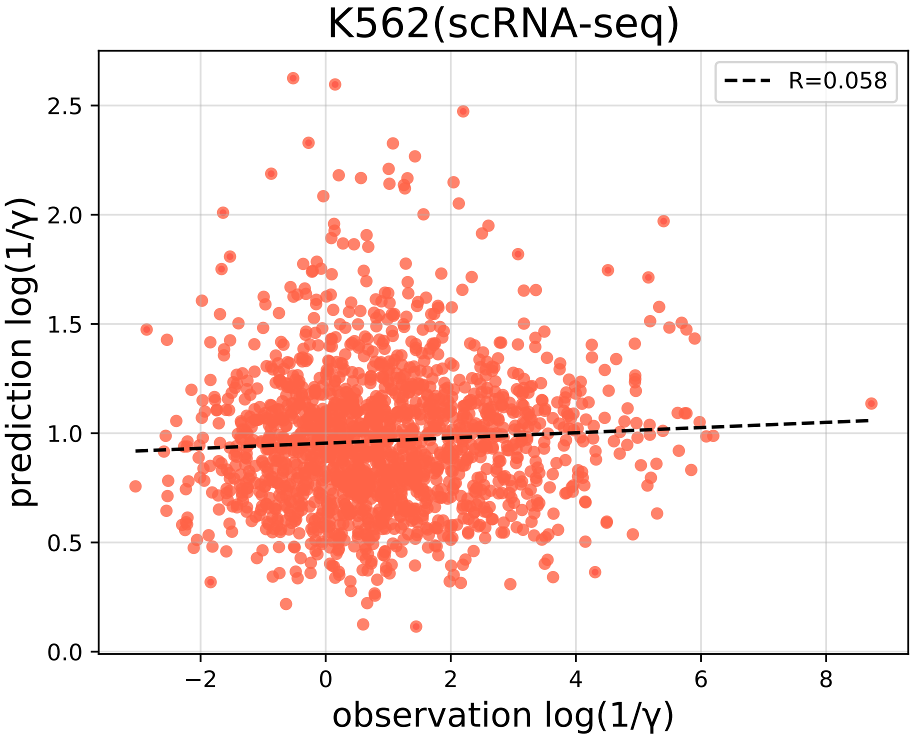


Figure S7: Poor prediction of splicing efficiency after permutation. Scatter plot of the relative splicing rate $(1/\gamma)$ between estimated by scVelo and predicted by 92 basic features, 96 octamer features and 120 RBP features after permuted across genes.

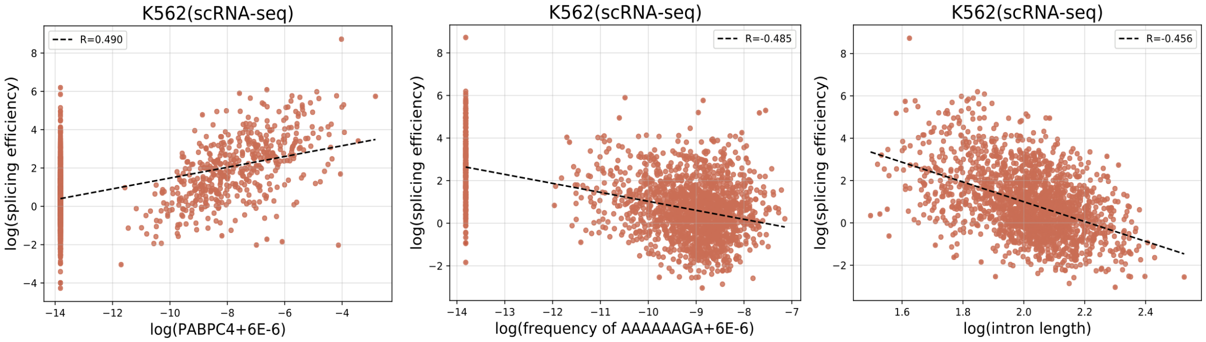


Figure S8: The scatter plot between relative splicing rate and the most important feature in each feature set for its prediction in K562 cell line: the RBP feature PABPC4 (left panel), the octamer AAAAAGA (middle panel) and the 92 basic feature intron length (right panel).

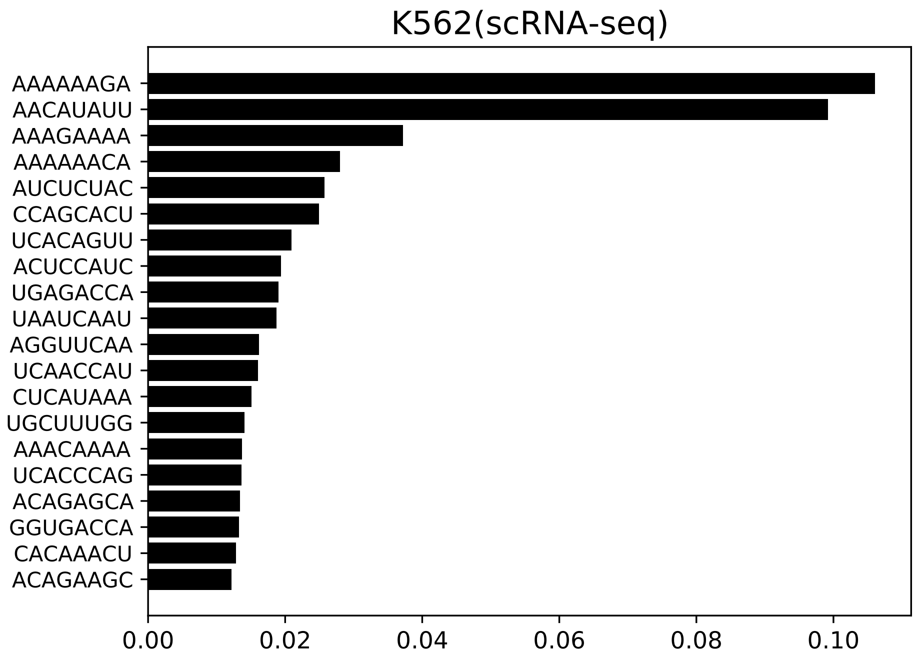


Figure S9: Top 20 important octamers feature to predict the relative splicing rate. Only the octamers feature set was applied for the prediction in a random forest regression. The importance index was shown on x-axis and the top 20 is presented.

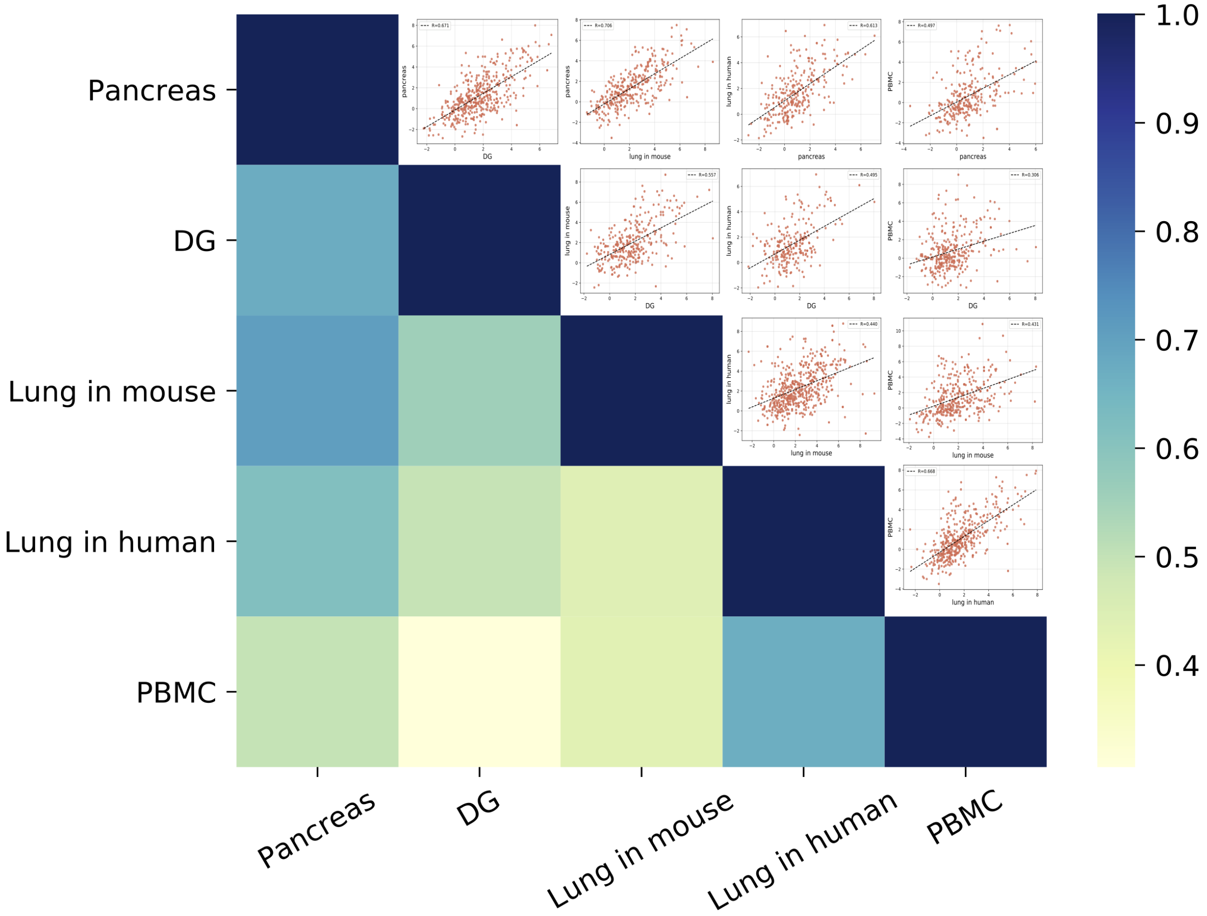


Figure S10: The pair-wise correlation of relative splicing efficiency across different tissues. The Pearson's R range from 0.306 to 0.706, indicated in the scatter plot and by the color in the heatmap plot.

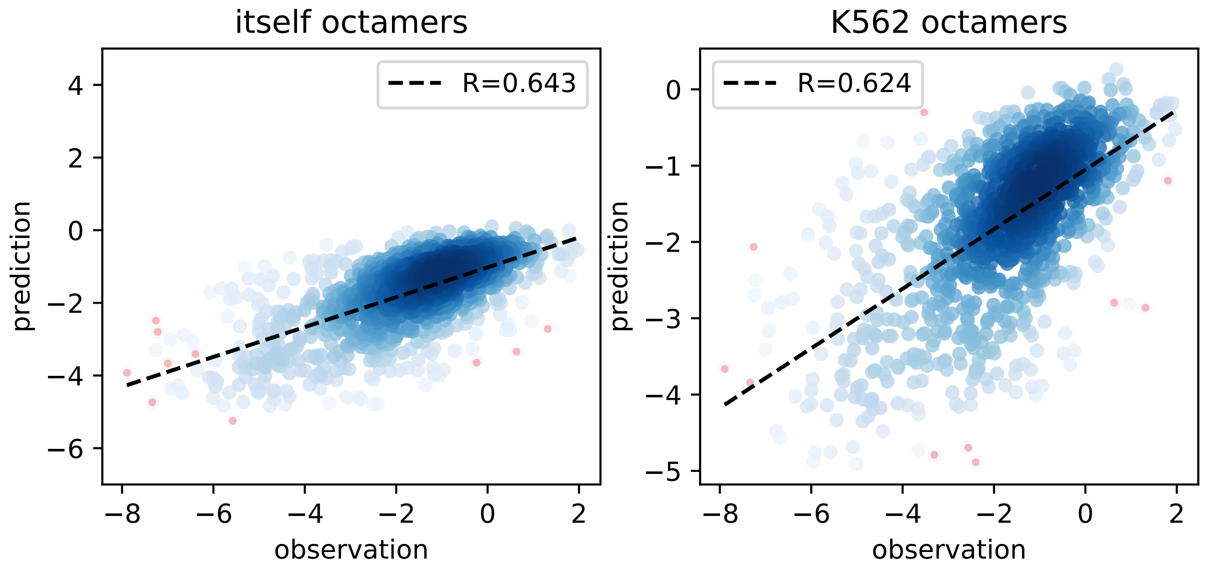


Figure S11: Selected octamers on K562 (scRNA-seq) are similarly predictive on other tissues. Scatter plot of $\log(1/\gamma)$ in human lung between the estimated by scVelo and the predicted by random forest regression with three feature sets (the 92 basic features set, the octamer features set and the RBP features set). Left panel: use the octamer features selected by the relative splicing rate in lung human dataset; Right panel: use the octamer features selected by the relative splicing rate in K562 (scRNA-seq) dataset.

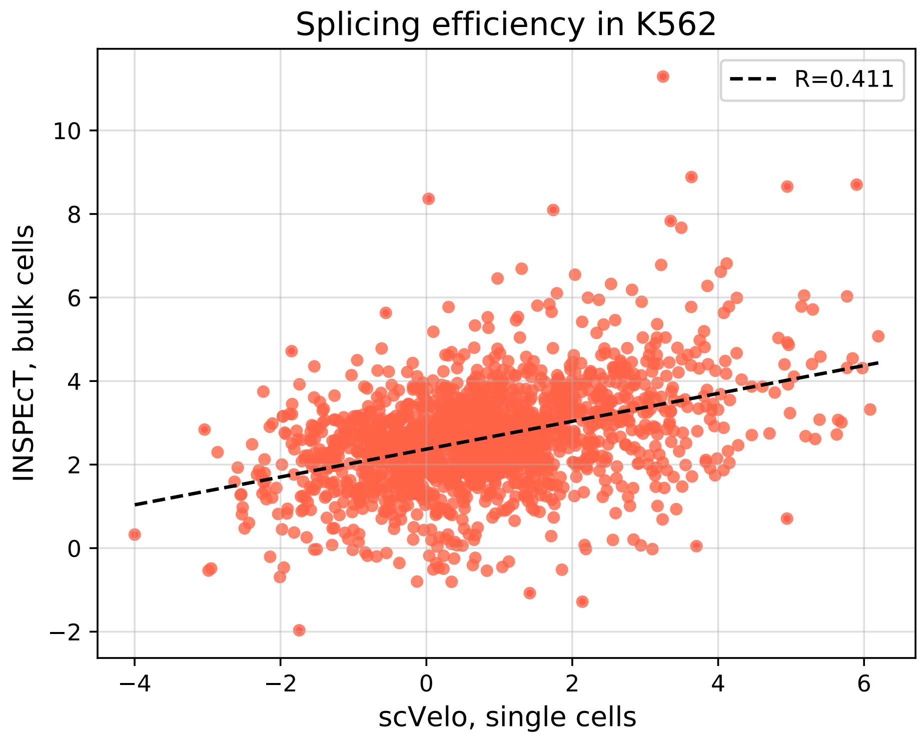


Figure S12: Scatter plot of the splicing efficiency estimates from K562. Bulk cells by INSPEcT- and single cell by scVelo.

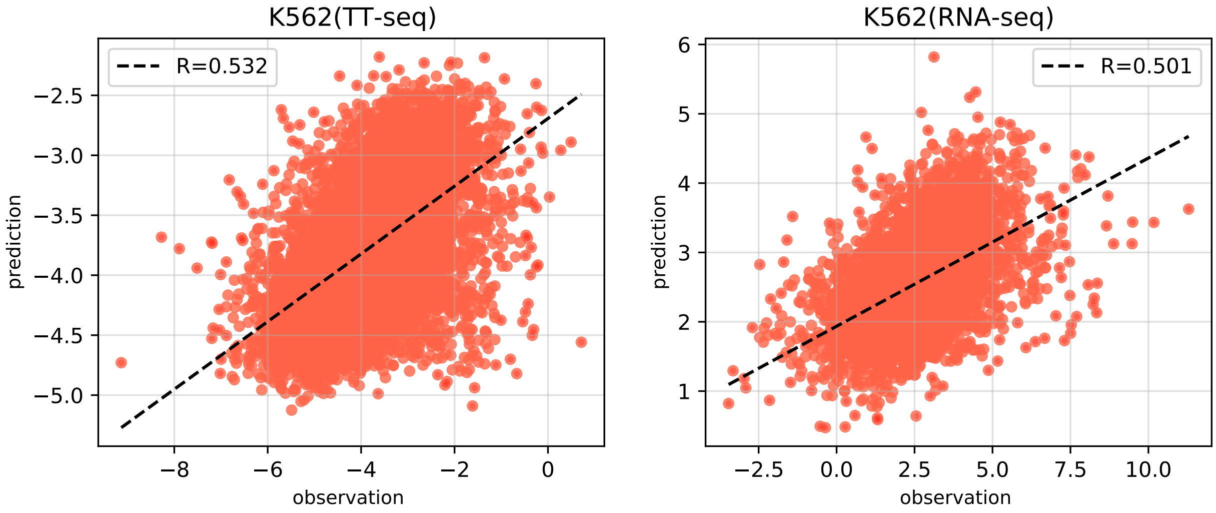


Figure S13: Scatter plot of the splicing efficiency estimated from different technique and predicted by three feature sets. The estimating splicing efficiency comes from the TT-seq (with metabolic label, left panel) and RNA-seq (without metabolic label, right panel).
